# A modular workflow for biomedical entity normalization across heterogeneous biological ontologies

**DOI:** 10.64898/2026.11.17.767229

**Authors:** Ana Amil, Monica Chagoyen

## Abstract

Understanding the biological mechanisms underlying rare diseases requires integrating information about phenotypes, cell types, anatomical structures, and molecular functions and processes. Although some of this knowledge is available in structured resources, additional relationships remain dispersed across biomedical literature. Extracting and integrating this information requires normalizing heterogeneous biological entities against specialized ontologies. This motivated the development of a common biomedical named entity normalization (NEN) workflow and the evaluation of its performance across different entity types and ontologies. We developed a modular NEN workflow combining exact matching, lexical and semantic candidate retrieval, and reranking, without requiring ontology-specific training. Candidate retrieval integrates BM25, fuzzy matching, character n-gram similarity, and SapBERT. We evaluated the workflow across the Human Phenotype Ontology (HPO), Cell Ontology (CL), Uberon, and Gene Ontology (GO) using multiple benchmarks, separately assessing candidate retrieval, reranking accuracy, and overall normalization accuracy. We also investigated factors associated with retrieval difficulty and compared the workflow with CellExLink, a specialized cell-type normalization system. Retrieval performance varied across ontologies and benchmarks, with lexical and semantic methods providing complementary candidate coverage in some settings. Increasing candidate-set size improved retrieval recall but did not necessarily improve overall normalization accuracy, as larger candidate sets could reduce reranking performance. Retrieval difficulty was associated with mention characteristics and ontology categories. Our evaluation identifies the performance characteristics and limitations of a common normalization workflow across heterogeneous biological ontologies, highlighting the importance of assessing candidate retrieval and selection separately. The workflow provides a methodological foundation for extracting and integrating biological information from rare disease literature, with potential applicability to other biomedical information-extraction tasks requiring normalization against these ontologies.

## Introduction

Understanding the biological mechanisms underlying rare diseases requires integrating information across different levels of biological organization, including phenotypes, molecular functions, biological processes, cellular components, cell types, and anatomical structures. Although established biomedical resources provide structured knowledge, such as disease–phenotype associations in the Human Phenotype Ontology (HPO; Gargano et al., 2024) and gene annotations in the Gene Ontology (GO; The Gene Ontology Consortium et al., 2026), additional relationships connecting these biological levels remain dispersed across the scientific literature. Systematically extracting and integrating this information requires identifying heterogeneous biological entities in text and mapping their mentions to standardized concepts in specialized ontologies, including HPO, GO, the Cell Ontology (CL; Tan et al., 2026), and Uberon, a cross-species anatomy ontology (Mungall et al., 2012). This mapping process, known as biomedical named entity normalization (NEN), or biomedical entity linking, is essential for transforming literature-derived information into structured, interoperable knowledge. Normalizing entities across these different ontologies presents a practical challenge for information-extraction applications that aim to integrate biological knowledge across multiple levels of organization.

Recent advances in biomedical language models have substantially improved entity normalization. Methods such as BioSyn (Sung et al., 2020) and SapBERT (Liu et al., 2020) have demonstrated the effectiveness of learning representations from biomedical terminology and synonym relationships, while other approaches have incorporated knowledge-base information and generative modeling (Yuan et al., 2022). These developments have provided methods applicable to different biomedical entity types and terminology resources. Nevertheless, normalization performance varies across entity types, terminologies, and evaluation benchmarks, as highlighted by the Biomedical Entity Linking Benchmark (BELB; Garda et al., 2023). Thus, the availability of broadly applicable normalization methods does not establish how well they perform across the particular biological ontologies required for a given information-extraction application.

Understanding these differences in performance requires examining the stages involved in normalization. Some mentions can be resolved through exact matching against ontology labels or synonyms, whereas others require methods capable of handling lexical and semantic variation. For these non-exact mentions, normalization typically involves candidate retrieval, which identifies plausible concepts, followed by candidate selection, which determines the most appropriate concept among them. Errors can arise either because the correct concept is not retrieved or because it is not selected despite being among the candidates. These stages also involve a potential trade-off: expanding candidate sets can improve retrieval recall while making the subsequent selection task more difficult. Evaluating these stages separately, with particular attention to mentions that cannot be resolved through exact matching, allows us to identify the main sources of normalization errors, characterize differences in difficulty across biological entity types and ontologies, and assess the limitations of the workflow for automated biological knowledge integration.

Motivated by the need to normalize heterogeneous biological information from rare disease literature, we developed a modular NEN workflow combining exact matching, lexical and semantic candidate retrieval, and candidate reranking. We adopted a common workflow rather than combining separate ontology-specific systems, aiming to simplify integration and maintenance and facilitate extension to additional ontologies. The workflow does not require ontology-specific training or architectural modifications, although ontology-adapted models can be incorporated when available. We evaluated its performance across HPO, CL, Uberon, and GO using available annotated corpora and complementary UMLS-based benchmarks (Bodenreider, 2004). We assessed exact matching and, for non-exact mentions, separately evaluated candidate retrieval, reranking accuracy conditional on successful retrieval, and overall normalization accuracy. We also investigated associations between retrieval difficulty and mention and ontology characteristics, and compared our workflow with CellExLink, a specialized cell-type normalization system. Together, these analyses characterize the workflow’s performance and limitations across the selected biological ontologies, providing a methodological foundation for rare disease literature mining and other biomedical information-extraction applications requiring these resources.

## Results

### Workflow implementation and evaluation datasets

We implemented a modular biomedical entity normalization workflow comprising three sequential stages: exact matching, candidate retrieval, and candidate reranking (Figure 1), following established approaches that combine lexical matching with candidate generation and ranking (Li et al., 2017; Sung et al., 2020, 2022). First, mentions are matched against ontology labels and synonyms, allowing exact matches to be directly assigned to ontology concepts. Mentions not resolved by exact matching are then processed using complementary lexical and semantic retrieval methods to identify candidate concepts. Finally, a reranking method selects the most appropriate concept from the retrieved candidates. The workflow combines training-free lexical methods with pretrained semantic representations and does not require ontology-specific model training or fine-tuning.

**Fig. 1.**
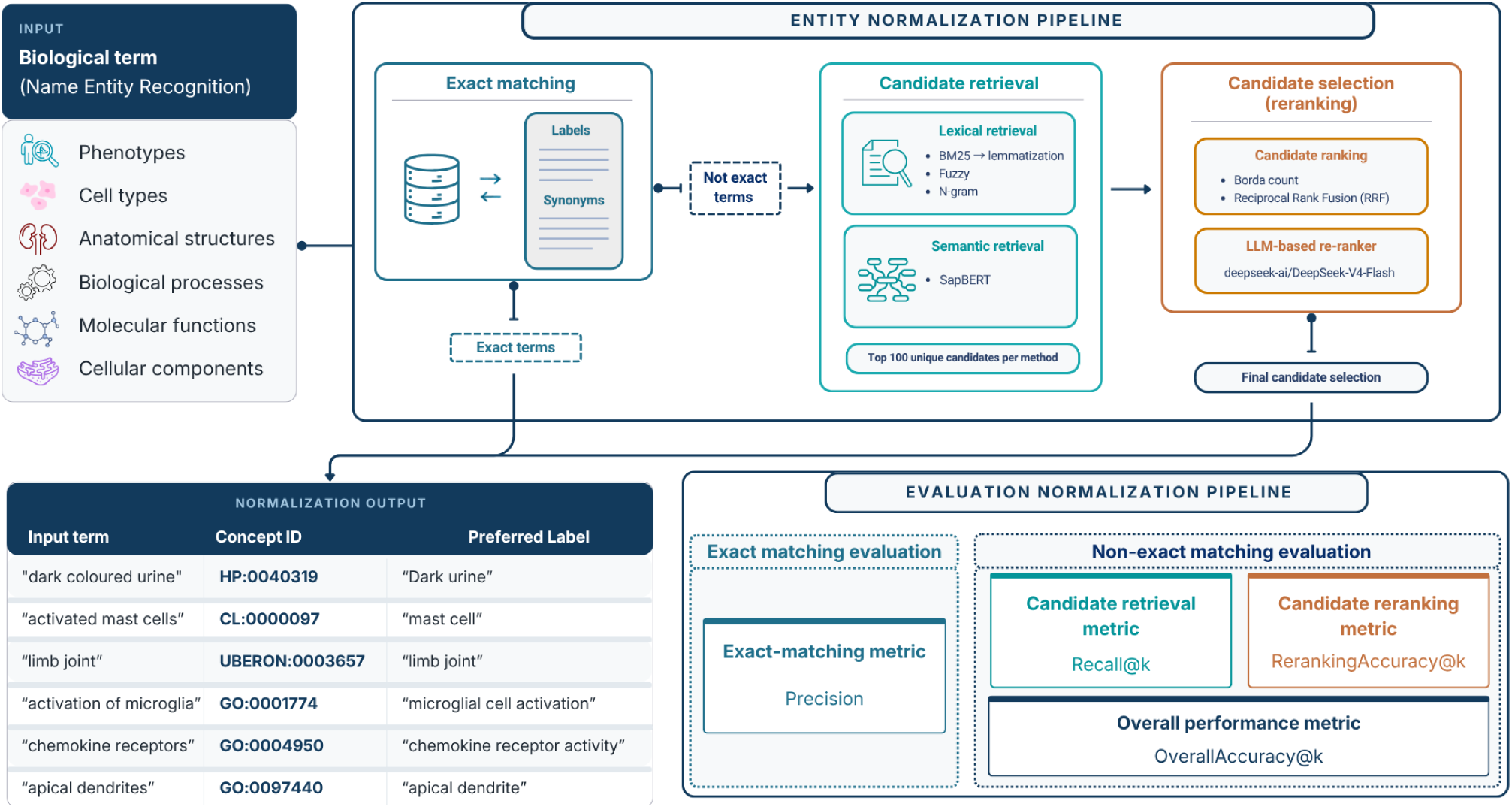
Overview of the biomedical entity normalization workflow and evaluation framework. Input mentions representing six types of biomedical entities are first matched against labels and synonyms in their corresponding target ontologies. Mentions without an exact match proceed to candidate retrieval, which combines lexical and semantic methods to identify and rank candidate concepts. Candidate reranking methods then selected the most appropriate concept from the retrieved candidates. The output consists of the input mention and the identifier of the selected concept in the corresponding ontology. Performance is evaluated at four levels: precision in exact-matching evaluation and candidate retrieval (Recall@k), candidate reranking (Reranking Accuracy@k), and overall normalization (Overall Accuracy@k) in non-exact matching evaluation.

We evaluated the workflow using four annotated corpora covering different biomedical entity types and terminology resources: MedMentions, CellLink, PhenoNormBench, and CRAFT (Bada et al., 2017; Mohan & Li, 2019; Rotenberg et al., 2026; Song et al., 2026). MedMentions contains annotations to UMLS concepts across multiple biomedical domains, which we grouped into semantic categories approximating the biological entity classes of interest (Methods). The other corpora provide annotations against specialized ontologies: CellLink for the Cell Ontology (CL), PhenoNormBench for the Human Phenotype Ontology (HPO), and CRAFT for the Gene Ontology (GO), CL, and Uberon. Together, these datasets provided complementary evaluation settings, including direct annotations against the target ontologies and broader UMLS-based annotations.

We first assessed the coverage and accuracy of exact matching and then evaluated candidate retrieval and reranking for mentions not resolved at that stage. By examining retrieval recall, reranking accuracy conditional on successful retrieval, and overall normalization accuracy, we distinguished errors arising from incomplete candidate retrieval from those occurring during final concept selection. This evaluation strategy allowed us to characterize the performance and limitations of the workflow across different biological entity types and terminology resources.

### Exact-match coverage and precision vary across ontologies and benchmarks

We first assessed the proportion of biomedical mentions that could be normalized through exact matching against the preferred labels and synonyms of the target ontologies. The proportion of non-exact mentions varied substantially across evaluation datasets and entity classes (Table 1). CellLink had the highest proportion of non-exact mentions (90.6%), followed by CRAFT (78.1%), PhenoNormBench (73.1%), and MedMentions (61.8%). Considerable differences were also observed between entity classes within the same dataset. For example, in MedMentions, the proportion of non-exact mentions ranged from 20.4% for anatomical entities to 73.7% for molecular functions. These differences indicate that the contribution of exact matching to overall normalization performance depends strongly on the evaluation corpus, entity class, and target ontology. They also highlight the importance of accounting for exact-match coverage when comparing normalization performance across benchmarks. Particularly striking differences were observed for anatomical entities: only 0.3% of CRAFT anatomical terms were non-exact matches, compared with 20.4% in MedMentions.

**Table 1.** Percentage of non-exact terms in each benchmark and entity type.

| **Entity type** | **MedMentions** | **CellLink** | **CRAFT** | **PhenoNormBench** | **FMA-Uberon** |
|---|---|---|---|---|---|
| Cell types | 71.5% | 90.6% | 80.6% | – | – |
| Phenotypes | 50.1% | – | – | 73.1% | – |
| Anatomical structures | 20.4% | – | 0.3% | – | 10.2% |
| Biological process | 66.9% | – | 70.3% | – | – |
| Molecular function | 73.7% | – | 92.5% | – | – |
| Cellular component | 59.0% | – | 75.4% | – | – |
| Overall | 61.8% | 90.6% | 78.1% | 73.1% | 10.2% |

The particularly high exact-match coverage of anatomical terms in CRAFT left only 19 non-exact mentions for evaluating retrieval performance against Uberon. To obtain a larger evaluation set, we constructed an additional dataset comprising 18,812 terms from the Foundational Model of Anatomy (FMA) that did not exactly match Uberon labels or synonyms and had corresponding Uberon concepts identified through ontology mappings (Methods). Unlike the annotated corpora, this dataset evaluates retrieval across anatomical terminologies rather than normalization of mentions occurring in biomedical text. We refer to this dataset as FMA–Uberon. The absolute numbers of unique entities in each benchmark are reported in Table S1.

We next evaluated the precision of exact-match assignments against the benchmark reference annotations (Table 2). Strict precision was generally high but varied across datasets, ranging from 0.770 for GO molecular functions in CRAFT to 0.997 for Uberon annotations in the same corpus. We additionally examined whether predicted concepts were ancestors or descendants of the reference concepts, using ontology relationships as alternative criteria in a post-hoc evaluation. Accepting hierarchically related concepts generally increased precision, particularly when descendants of the reference concepts were included. For example, precision for GO molecular functions in CRAFT increased from 0.770 to 0.846 when both ancestor and descendant relationships were accepted. These differences suggest that some apparent normalization errors reflect differences in ontological specificity rather than assignments to unrelated concepts. Detailed precision results for the MedMentions benchmark are reported in Table S2.

**Table 2.**
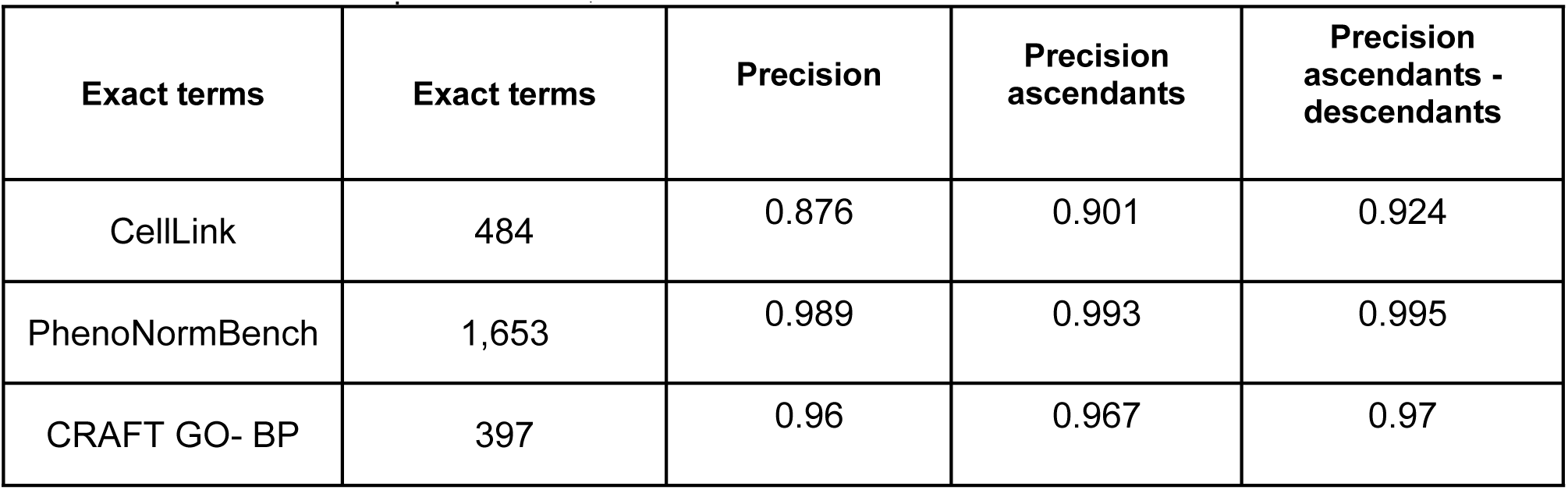
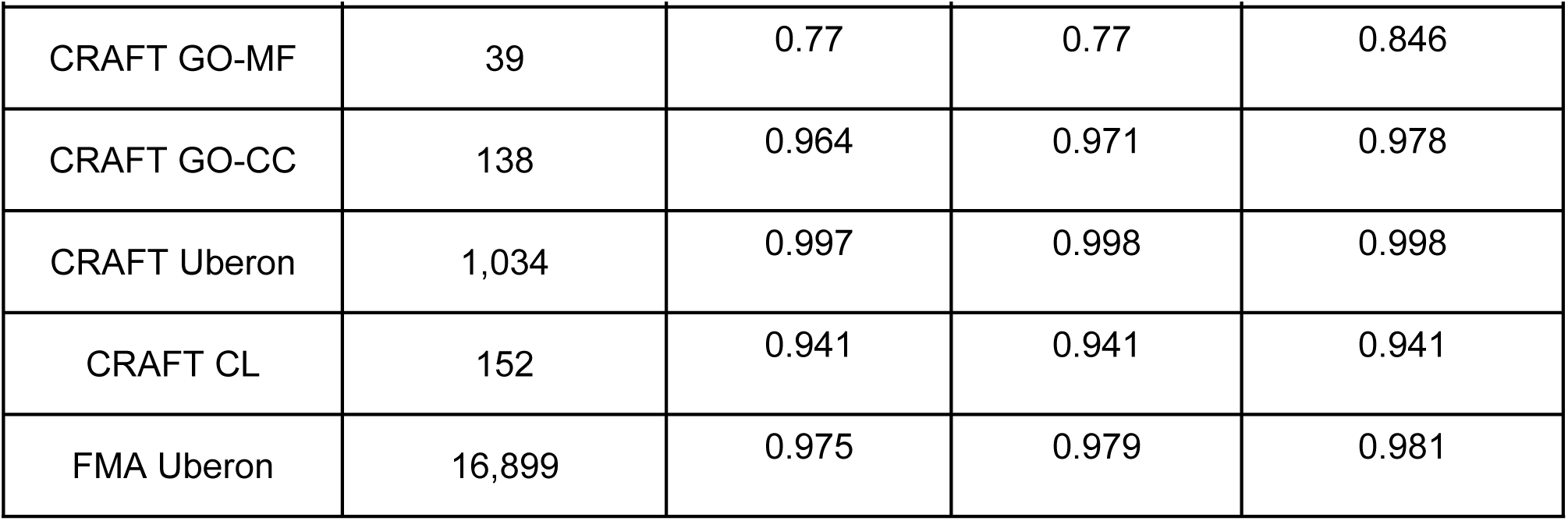
Precision of exact terms. Number of distinct exact-match terms and exact-match precision including also hierarchical relationships in CellLink, PhenoNormBench and CRAFT benchmarks.

To better understand discrepancies between exact-match predictions and benchmark annotations, we manually inspected all predictions classified as incorrect under strict evaluation in CellLink (60 cases), CRAFT GO molecular functions (9 cases), and CRAFT GO cellular components (5 cases). This analysis identified several types of disagreement:

- **Potential benchmark annotation errors.** In CRAFT, the mention *PKA* was annotated as *AMP-activated protein kinase activity*, whereas exact matching selected *cAMP-dependent protein kinase activity*, consistent with the conventional meaning of the abbreviation.
- **Ambiguous abbreviations.** In CellLink, *MC* was associated with different reference concepts, including *mast cell*, *macrophage*, and *monocyte*, whereas exact matching selected *Merkel cell*. This illustrates the difficulty of resolving ambiguous abbreviations through lexical matching alone.
- **Differences in annotation specificity.** In CRAFT, *PFK* was annotated as *phosphofructokinase activity*, whereas exact matching selected the more specific *6-phosphofructokinase activity*. Similarly, *nerve ending* was annotated as *neuron projection terminus*, whereas exact matching selected *axon terminus*, for which *nerve ending* is listed as an exact synonym in GO. In both cases, the predicted concept was more specific than the reference annotation.
- **Generic or underspecified mentions.** In the cellular component subset of CRAFT, *ribbon* was annotated as *synaptic ribbon*, whereas exact matching selected *cytoskeleton of presynaptic active zone*, illustrating the ambiguity of generic terms without sufficient contextual information.

Overall, exact matching resolved a variable proportion of mentions with generally high precision. Importantly, the analysis also revealed limitations of strict agreement with benchmark annotations as a measure of normalization correctness. Although exact lexical matching does not guarantee a correct assignment, some discrepancies may reflect ambiguous terminology, differences in annotation specificity, or potentially incorrect reference annotations rather than normalization errors. Consequently, imperfections in benchmark annotations may impose a practical ceiling on the accuracy measured against reference identifiers, even for otherwise appropriate predictions. The hierarchy-aware evaluation further illustrates that strict identifier agreement may not fully capture the biological or ontological appropriateness of a normalization. These considerations are relevant when interpreting the performance of the subsequent candidate retrieval and reranking stages, particularly for non-exact mentions where determining the appropriate concept may be more challenging.

### Retrieval methods show variable performance across biomedical entity types and ontologies

Mentions that do not exactly match ontology labels or synonyms require lexical or semantic generalization. We therefore restricted the evaluation of candidate retrieval and reranking to non-exact mentions, allowing us to assess performance beyond direct terminology matching. Candidate retrieval was evaluated using three lexical methods (BM25, fuzzy matching, and character n-gram similarity) and one semantic method (SapBERT), measuring the proportion of reference concepts retrieved among the top-ranked candidates (Recall@k; Methods).

#### Retrieval performance across biomedical entity types in MedMentions

We first evaluated retrieval performance across different biomedical entity types using MedMentions, a corpus annotated with UMLS concepts. To approximate the biological entity classes of interest, we grouped mentions according to selected UMLS semantic types corresponding to cell types, anatomical entities, phenotypes, biological processes, molecular functions, and cellular components (Methods).

We compared Recall@k for each retrieval method individually and for their combined candidate sets (Figure 2). For the combined approach, the top-k candidates from each method were merged and duplicates removed. Because candidate overlap resulted in fewer than 4k unique concepts, the combined candidate set size was represented by the median number of unique candidates across mentions.

**Fig. 2.**
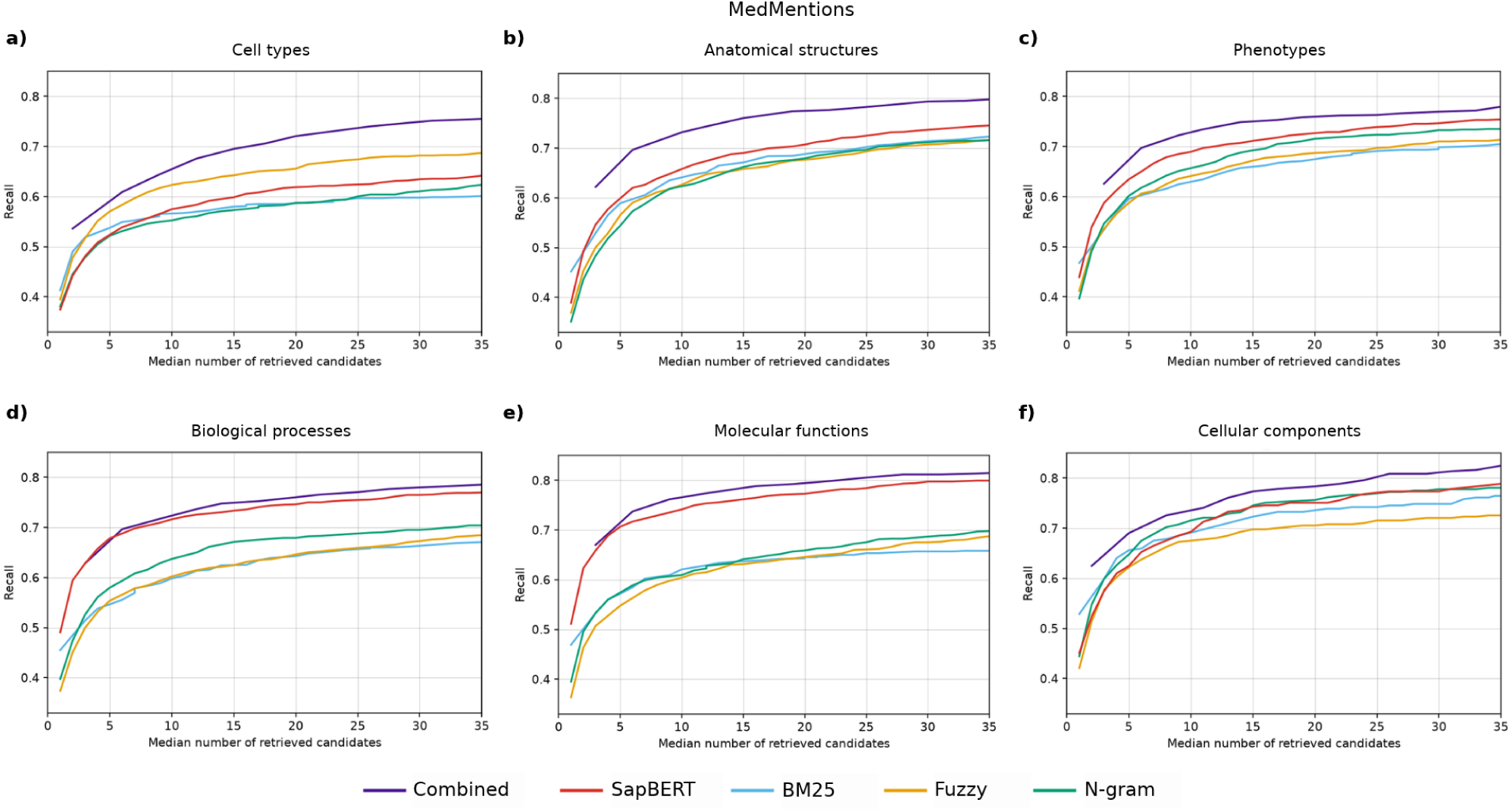
Candidate retrieval performance across biomedical entity types in MedMentions. Recall@k as a function of the number of retrieved candidates for BM25, fuzzy matching, character n-gram similarity, SapBERT, and their combined candidate sets. Results are shown separately for (a) cell types, (b) anatomical entities, (c) phenotypes, (d) biological processes, (e) molecular functions, and (f) cellular components. For the combined approach, the top-k candidates from each method were merged and deduplicated; the x-axis represents the median number of unique candidates across mentions. Only mentions without an exact match to the target terminology were included.

Retrieval performance varied across entity types and methods. Recall generally increased rapidly for the first few candidates, followed by progressively smaller gains as the candidate set expanded. However, the rate of improvement differed across entity types, with some curves approaching a plateau at relatively small candidate sets and others continuing to increase up to the maximum evaluated size. To illustrate the range of retrieval performance across methods and entity types, we compared recall at two reference points. Recall@1 ranged from 0.351 to 0.529, with the lowest values observed for cell types and anatomical entities. At k = 25, recall ranged from 0.597 to 0.805, with the highest value achieved by SapBERT for molecular functions.

The three lexical methods generally showed comparable performance within each entity type, although fuzzy matching performed better for cell types and character n-gram matching for biological processes. SapBERT consistently outperformed the lexical methods for biological processes and molecular functions.

Combining candidates from all four methods improved recall over individual methods for cell types, anatomical entities, and phenotypes, with a smaller improvement for cellular components. For biological processes and molecular functions, the combined approach achieved recall comparable to SapBERT alone at similar candidate-set sizes. These findings indicate that combining lexical and semantic retrieval methods can improve candidate recall for some biomedical entity types, whereas for others, the combined approach provides little additional recall over SapBERT alone.

#### Retrieval performance in ontology specific benchmarks

We next evaluated candidate retrieval against specialized biomedical ontologies using CellLink for CL, PhenoNormBench for HPO, CRAFT for GO biological processes, molecular functions, and cellular components, and FMA–Uberon for anatomical concepts (Figure 3). For CL, we selected CellLink because it provided a substantially larger set of non-exact mentions than CRAFT (4,672 versus 637).

**Fig. 3.**
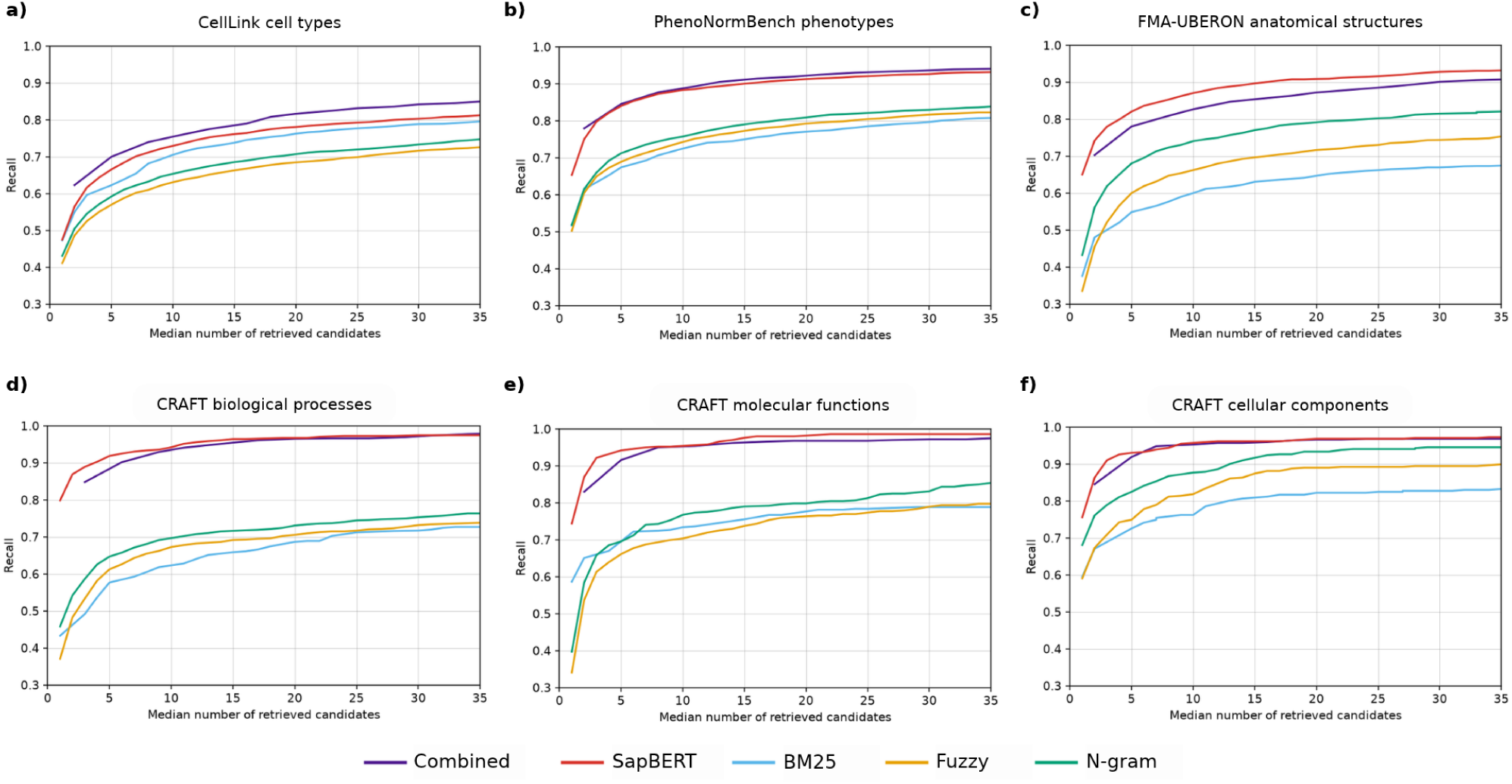
Candidate retrieval performance across specialized biomedical ontologies. Recall@k for the evaluated retrieval methods across six ontology-specific datasets and entity classes: (a) Cell Ontology (CL) in CellLink, (b) Human Phenotype Ontology (HPO) in PhenoNormBench, (c) Uberon in FMA–UBERON, and Gene Ontology (GO) annotations in CRAFT for (d) biological processes, (e) molecular functions, and (f) cellular components.

SapBERT consistently outperformed the individual lexical retrieval methods across all evaluated ontologies, with particularly pronounced differences for phenotypes, biological processes, and molecular functions. In CellLink and PhenoNormBench, SapBERT achieved Recall@3 values of 0.616 and 0.798, respectively, increasing to 0.793 and 0.921 at k = 25. The relative performance of lexical methods also varied across entity classes. Character n-gram similarity performed particularly well for anatomical concepts and cellular components, whereas BM25 achieved higher recall for cell types. In contrast, the three lexical methods showed broadly comparable performance for phenotypes, biological processes, and molecular functions.

Combining candidates from all four retrieval methods improved recall over SapBERT alone in CellLink, whereas the combined approach provided little or no additional benefit in the other evaluations. In FMA–Uberon, SapBERT outperformed both the individual lexical methods and their combined candidate sets. Despite the improvement obtained by combining methods, CellLink exhibited substantially lower recall than the other benchmarks, even at the largest evaluated candidate-set size. These results indicate that retrieving reference concepts was particularly challenging for cell-type mentions in CellLink, motivating further investigation of factors associated with retrieval performance and comparison with a specialized cell-type normalization system.

### Candidate retrieval and concept discrimination constitute distinct normalization bottlenecks

We next evaluated the ability of different reranking methods to select the correct concept from retrieved candidates, independently of candidate retrieval performance. We compared an LLM-based reranker with two rank-aggregation methods, Borda count and reciprocal rank fusion (RRF). Reranking accuracy was calculated among mentions for which the reference concept was successfully retrieved, whereas overall normalization accuracy was calculated across all evaluated mentions, thereby accounting for both candidate retrieval and final concept selection (Methods).

We focused on CellLink and PhenoNormBench, which provided the largest sets of non-exact mentions. Based on the preceding retrieval results, we selected two candidate-generation strategies: SapBERT, which achieved the highest recall among individual methods, and the combination of all four retrieval methods, which could provide additional candidate coverage through complementary retrieval approaches. Comparing these strategies allowed us to assess whether broader candidate retrieval improved final normalization accuracy or introduced additional difficulties for concept discrimination. To ensure comparable candidate-set sizes, we used the median number of unique candidates per mention after merging and deduplication. The resulting reranking and overall normalization accuracies across candidate-set sizes are shown in Figure 4. The performance representations for the remaining benchmarks are presented in the Supplementary Material (Figure S1).

**Fig. 4.**
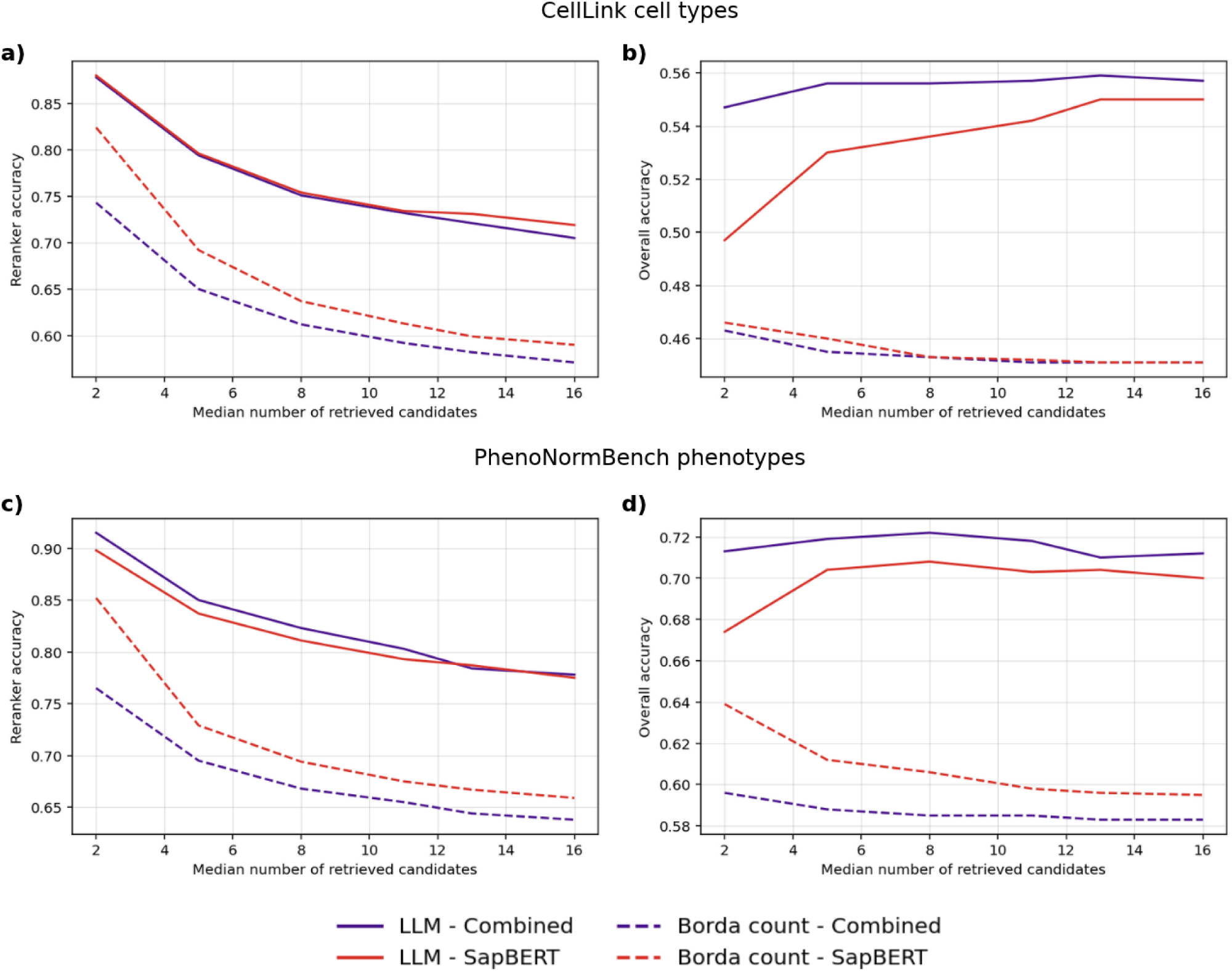
Candidate selection performance across CellLink and PhenoNormBench. Reranker and Overall Accuracy@k across candidate-set sizes. These metrics are calculated using LLM-based and Borda count rerankers in two retrieval scenarios: combined candidates, consisting of candidates retrieved by lexical and SapBERT methods and only SapBERT candidates. Results of each metric and benchmark are shown separately: (a) Reranker accuracy@k in CellLink benchmark, (b) Overall accuracy@k in CellLink benchmark, (c) Reranker accuracy@k in PhenoNormBench benchmark and (d) Overall accuracy@k in PhenoNormBench benchmark.

The LLM-based reranker consistently outperformed Borda and RRF in both benchmarks. At approximately two retrieved candidates, the LLM achieved conditional reranking accuracies of 0.915 in PhenoNormBench and 0.878 in CellLink using combined retrieval, compared with 0.765 and 0.743 for Borda, and 0.763 and 0.738 for RRF, respectively. Borda and RRF showed nearly identical performance across these comparisons.

Increasing candidate-set size produced competing effects on normalization performance. Although larger candidate sets improved the probability of retrieving the reference concept, they also increased the difficulty of selecting the correct candidate. Consequently, overall normalization accuracy did not increase monotonically with candidate-set size. For the LLM with combined retrieval, maximum overall accuracy was 0.722 in PhenoNormBench at approximately eight candidates and 0.559 in CellLink at approximately 13 candidates. With SapBERT-only retrieval, the corresponding maxima were 0.708 and 0.550, respectively. These results indicate that the candidate-set size maximizing overall accuracy differs between benchmarks.

We additionally assessed normalization performance using ontology relationships in a post-hoc evaluation. Numerical results for all the selected evaluation benchmarks are presented in supplementary metrics Table S3-S8. Predictions were alternatively accepted when they corresponded to ancestors of the reference concept or, under a more permissive criterion, to either ancestors or descendants. Both criteria increased measured accuracy, indicating that some disagreements under strict identifier matching involve differences in ontological specificity. For example, acinar cells was annotated as acinar cell of salivary gland in CellLink, whereas the workflow selected the more general acinar cell. However, hierarchical relatedness alone does not establish that a prediction is appropriate in its textual context. These discrepancies are examined further in the subsequent error analysis.

Together, these results show that candidate retrieval and concept discrimination impose distinct limitations on normalization performance. The LLM provided more accurate candidate selection than rank aggregation, but the benefit of increasing candidate coverage depended on the reranker’s ability to discriminate among the additional candidates.

### Factors associated with candidate retrieval performance in CellLink

Given the comparatively low candidate retrieval recall observed in CellLink and its large number of non-exact mentions, we selected this benchmark for a more detailed investigation of factors associated with retrieval performance. We first examined whether the rank assigned to the reference concept was associated with mention length, measured by the number of characters and tokens, and with the number of synonyms associated with the reference concept in the Cell Ontology (CL).

All four retrieval methods showed statistically significant negative correlations between reference concept rank and mention length, indicating that shorter mentions tended to receive poorer rankings (Table 3). The strongest associations were observed for BM25 (Spearman’s ρ = −0.360 for character count and −0.313 for token count), whereas the corresponding correlations were weaker for the other methods (ρ ranging from −0.269 to −0.136). The number of synonyms associated with the reference concept was also negatively correlated with retrieval rank across all methods, although these associations were relatively weak (ρ ranging from −0.181 to −0.124). Thus, both mention length and synonym coverage were associated with retrieval performance, with mention length showing a more pronounced relationship, particularly for BM25.

**Table 3.**
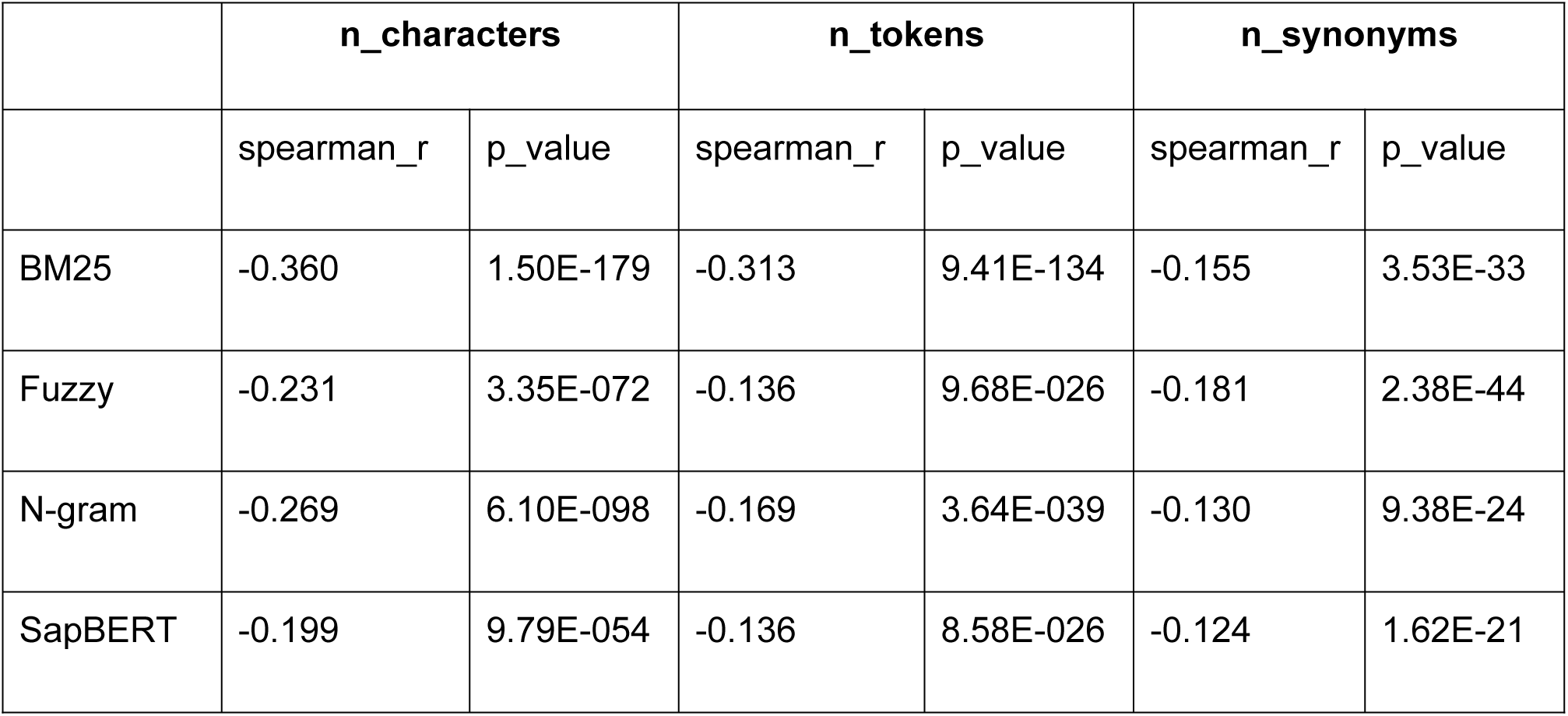
Spearman’s rank correlation coefficient between retrieval rank and query terms characteristics per each retrieval method.

|  | **n_characters** |  | **n_tokens** |  | **n_synonyms** |  |
|  | spearman_r | p_value | spearman_r | p_value | spearman_r | p_value |
|---|---|---|---|---|---|---|
| BM25 | -0.360 | 1.50E-179 | -0.313 | 9.41E-134 | -0.155 | 3.53E-33 |
| Fuzzy | -0.231 | 3.35E-072 | -0.136 | 9.68E-026 | -0.181 | 2.38E-44 |
| N-gram | -0.269 | 6.10E-098 | -0.169 | 3.64E-039 | -0.130 | 9.38E-24 |
| SapBERT | -0.199 | 9.79E-054 | -0.136 | 8.58E-026 | -0.124 | 1.62E-21 |

We next investigated whether retrieval performance differed between mentions referring to a single cell type (5,029) and mentions containing lists of cell types (865) e.g. “acinar, ductal and myoepithelial cells”. For list-based mentions, retrieval was considered successful if any of the annotated reference concepts was retrieved, rather than requiring all of them. Despite this permissive criterion, the distributions of reference concept ranks differed significantly between the two groups for all four retrieval methods (two-sided Kolmogorov–Smirnov test; Table 4). List-based mentions generally received poorer rankings, with median ranks increasing from 7 to 8 for BM25, from 6 to 7 for fuzzy matching, from 6 to 9 for character n-gram similarity, and from 3 to 5 for SapBERT. Although SapBERT achieved the lowest median ranks in both groups, its performance was also affected by mention composition. These results suggest that mentions containing multiple cell-type expressions present additional retrieval difficulties, even when recovering only one of the annotated concepts is sufficient.

**Table 4.** Two-sided Kolmogorov-Smirnov (KS) test to determine distribution of ranks between single term queries and list of terms.

|  | **Median rank single term** | **Median rank list of terms** | **KS_statistic** | **p_value** |
|---|---|---|---|---|
| BM25 | 7 | 8 | 0.186 | 6.267E-23 |
| Fuzzy | 6 | 7 | 0.117 | 2.499E-09 |
| N-gram | 6 | 9 | 0.133 | 7.644E-12 |
| SapBERT | 3 | 5 | 0.166 | 2.813E-18 |

Finally, we examined whether retrieval performance varied across hierarchical groups of the Cell Ontology. For each parent concept, we identified all reference concepts belonging to its transitive descendant hierarchy, including the parent itself. We then compared their retrieval rank distributions with those of concepts outside that hierarchy using the two-sided Kolmogorov–Smirnov test (Figure S2). Significant differences were identified for several hierarchical groups, indicating that retrieval performance was not uniform across cell-type categories.

We examined SapBERT in particular to investigate whether the semantic retrieval method showed patterns distinct from those of the lexical approaches. Two hierarchical groups showed significant differences specific to SapBERT: professional antigen presenting cell and supporting cell, whose associated concepts tended to receive poorer rankings. Other differences were shared with lexical methods. For example, SapBERT and BM25 showed poorer rankings for phagocyte, stuff accumulating cell, and stromal cell, whereas SapBERT and fuzzy matching showed better rankings for hematopoietic precursor cell, and SapBERT and character n-gram similarity showed poorer rankings for sensory epithelial cell. Additional differences observed among the lexical methods, but not SapBERT, involved stem cell, endo-epithelial cell, and neuron associated cell.

Overall, these analyses revealed associations between retrieval performance and mention length, synonym coverage, mention composition, and ontology-defined cell-type groups. The overlap in hierarchical patterns across semantic and lexical retrieval methods suggests that at least some differences may reflect characteristics of the mentions or reference concepts associated with particular cell-type groups, rather than properties specific to semantic retrieval. However, the present analyses do not establish the independent contributions of these factors.

#### Comparison with a specialized cell-type normalization system

We finally compared our general-purpose normalization framework with CellExLink (Nabijiang & Shahriyari, 2026), a specialized cell-type normalization system providing an implementation and evaluation data suitable for direct comparison. We used the CellExLink test subset of CellLink to evaluate both candidate retrieval and overall normalization accuracy.

We first compared the candidate retrieval performance of SapBERT with SapBERT-cell, a version fine-tuned by the CellExLink authors for cell-type normalization through supervised contrastive learning using Cell Ontology terminology and annotated cell-type mentions. SapBERT-cell consistently outperformed SapBERT across the evaluated range of candidate-set sizes, increasing Recall@1 from 0.521 to 0.594 (7.3 percentage points). The improvement was somewhat more pronounced for smaller candidate sets but remained evident at larger values of *k* (Figure 5a). We then evaluated SapBERT-cell alongside the three lexical retrieval methods and their combined candidate sets (Figure 5b).

**Fig 5.**
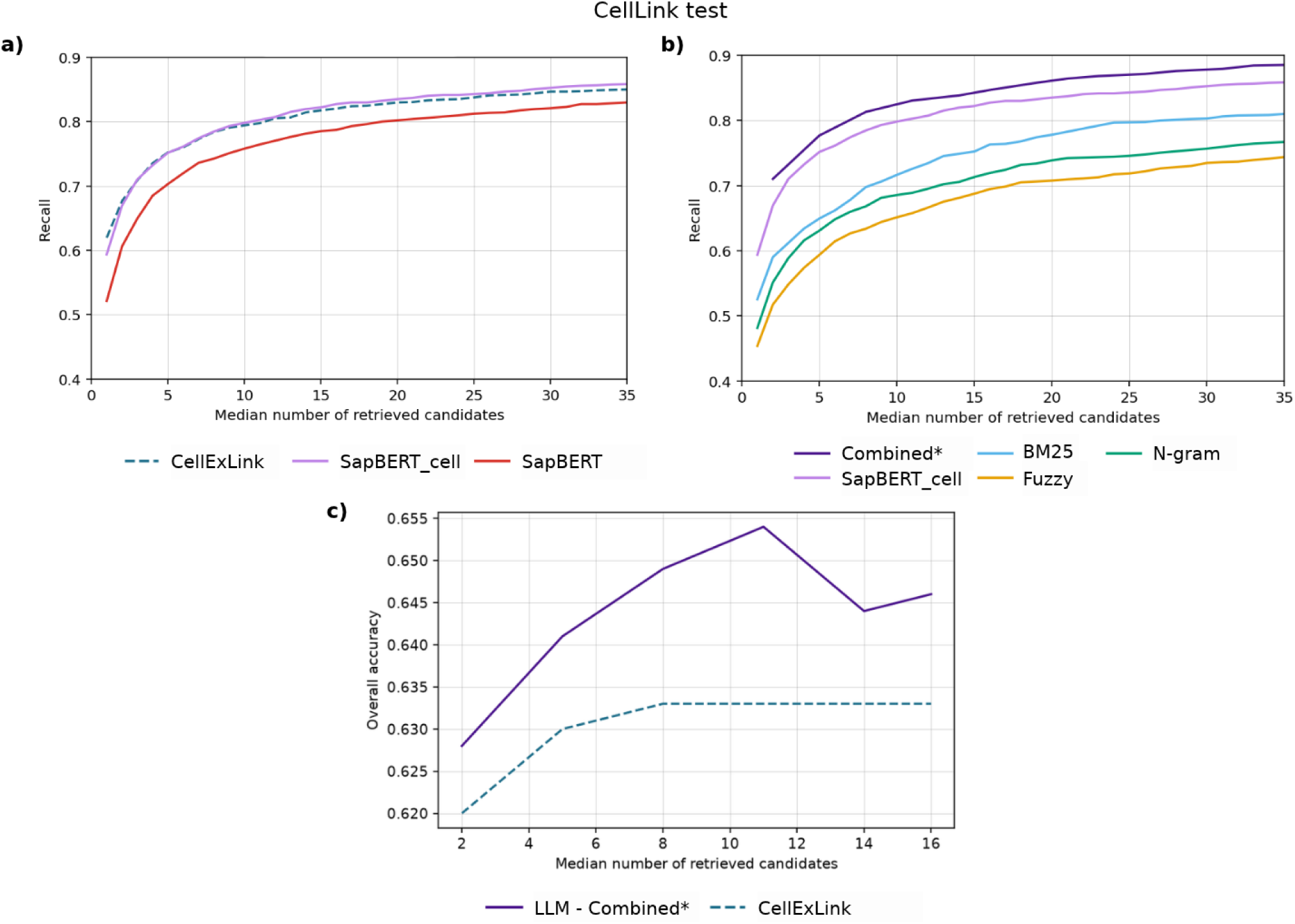
Candidate retrieval and reranking performance across CellLink test benchmark. (a) Recall@k for CellExLink, SapBERT_cell and SapBERT retrieval methods, (b) Recall@k of the lexical retrieval methods, SapBERT_cell and both types of retrieval methods combined, and (c) Overall Accuracy@k of two retrieval and reranking scenarios: LLM-based reranking across candidate-set sizes in combined candidates, consisting of candidates retrieved by lexical and SapBERT_cell methods, and candidates retrieved and reranked by CellExLink pipeline.

To investigate whether our general-purpose normalization framework could achieve comparable performance to a specialized system when incorporating an ontology-adapted retrieval model, we compared our pipeline with CellExLink on the same test subset. Our pipeline combined candidates retrieved by SapBERT-cell and the three lexical methods, followed by LLM-based reranking, whereas CellExLink used its own retrieval and trained reranking components. Both pipelines were evaluated using comparable median candidate-set sizes, ranging from approximately 2 to 16 candidates.

The two pipelines achieved similar overall normalization accuracy across the evaluated candidate-set sizes (Figure 5c). Our framework reached a maximum accuracy of 0.654 at a median of 11 candidates, compared with 0.633 for CellExLink, with differences ranging from 0.008 to 0.021 across the evaluated candidate-set sizes.

These results indicate that our general-purpose normalization framework can incorporate an ontology-adapted semantic retrieval model and achieve overall performance comparable to that of a specialized cell-type normalization system employing a trained reranker. Notably, CellExLink’s candidate retrieval recall was similar to that obtained with SapBERT-cell alone, suggesting that differences in overall normalization accuracy were not primarily driven by differences in candidate retrieval recall. Nevertheless, because the two pipelines do not necessarily operate on identical candidate sets, their comparison does not provide a direct assessment of the relative performance of the reranking methods. Normalization performance comparison between combined candidates and CellExLink candidate retrieval and reranking using ontology relationships in a post-hoc evaluation are presented in supplementary metrics Table S9.

## Discussion

Integrating biological knowledge from rare disease literature requires identifying and normalizing heterogeneous entities, including phenotypes, molecular functions, biological processes, cellular components, cell types, and anatomical structures. These entities are represented in different specialized ontologies, creating a practical need for normalization workflows that can operate across terminology resources without requiring separate models for each entity type. Motivated by this requirement, we developed and evaluated a modular workflow combining exact matching, lexical and semantic candidate retrieval, and candidate reranking. Our results characterize its performance across different biological ontologies and benchmarks, revealing substantial variation in retrieval difficulty and highlighting the complementary roles of candidate retrieval and selection in determining normalization accuracy.

### Normalization across heterogeneous biological ontologies

The workflow was designed to provide a common normalization approach across the biological ontologies required for our intended information-extraction application. Its general-purpose character lies in its ability to operate across different terminology resources without requiring ontology-specific training or architectural modifications, rather than in the development of new normalization algorithms. This does not imply that its components are independent of existing biomedical terminology resources. In particular, SapBERT was pretrained using UMLS synonym relationships, which include terminology from some of the ontologies evaluated here.

Candidate retrieval performance varied considerably across benchmarks, with SapBERT generally providing the strongest individual retrieval strategy. Combining semantic and lexical methods improved candidate coverage in some settings, particularly CellLink and several MedMentions entity groups, but provided little additional benefit in others. These findings indicate that lexical and semantic retrieval can provide complementary coverage, although their combination does not necessarily improve recall. The effectiveness of each strategy depends on the entity types, terminology resources, and evaluation datasets involved. Consequently, a common normalization workflow does not imply a single retrieval configuration that performs optimally across all ontologies.

The CellLink analyses provided further insight into factors associated with retrieval difficulty. Retrieval performance varied with mention length, synonym coverage, mention composition, and ontology-defined cell-type categories. Shorter mentions and mentions containing lists of cell types tended to receive poorer rankings, while some differences among cell-type categories were observed across both lexical and semantic methods. These findings suggest that normalization difficulty reflects not only the retrieval method but also characteristics of the mentions and reference terminology. However, these associations are exploratory and do not establish the independent contribution of individual factors.

### Candidate retrieval and selection as complementary challenges

An important finding of this study is that candidate retrieval and concept selection impose distinct limitations on normalization accuracy. Retrieving the correct concept is necessary but insufficient for successful normalization: the correct concept may be included among the candidates but not selected by the reranking method. Although increasing candidate-set size generally improved retrieval recall, it did not consistently improve overall normalization accuracy, as larger candidate sets could make concept selection more difficult. This trade-off emphasizes the importance of evaluating retrieval and reranking jointly. In particular, improving the ranking of correct concepts within relatively small candidate sets may be more beneficial than simply expanding candidate coverage when the selection method has difficulty distinguishing closely related concepts. Evaluating retrieval recall and reranking accuracy separately therefore provides information that would be obscured by considering overall normalization accuracy alone.

Among the evaluated candidate selection strategies, LLM-based reranking achieved the highest overall normalization accuracy in the principal benchmarks. It also benefited from combining candidates retrieved through different methods, whereas conventional rank-based approaches generally performed better with SapBERT candidates alone. These results suggest that LLM-based reranking can make effective use of heterogeneous candidate sets without ontology-specific training, although its performance remains dependent on the quality of candidate retrieval.

The comparison with CellExLink illustrates how ontology-specific adaptation can complement a common normalization workflow. SapBERT-cell, fine-tuned by the CellExLink authors using Cell Ontology terminology and annotated cell-type mentions, consistently improved candidate retrieval relative to pretrained SapBERT across the evaluated candidate-set sizes, with larger gains for smaller sets. Incorporating this model into our workflow, together with lexical retrieval and LLM-based reranking, achieved overall normalization accuracy comparable to that of the specialized CellExLink pipeline.

### Evaluation considerations and limitations

Several aspects of the evaluation should be considered when interpreting these results. First, the benchmarks differed in their target terminology resources, annotation conventions, and proportions of mentions that could be resolved through exact matching. Since exact matches can often be normalized through direct terminology lookup, their prevalence can substantially influence overall normalization accuracy. Our separate evaluation of non-exact mentions provides a more focused assessment of candidate retrieval and selection, although differences in benchmark composition still limit direct comparisons across ontologies. The evaluation also included both corpora annotated directly against the target ontologies and UMLS-based benchmarks. The latter provide complementary evidence across biomedical entity categories but do not necessarily reproduce the terminology scope or annotation characteristics of HPO, CL, Uberon, and GO. Consequently, performance observed in UMLS-based evaluations should not be interpreted as directly equivalent to normalization against the corresponding specialized ontologies.

Second, reference annotations do not always provide an unambiguous measure of normalization correctness. Some discrepancies may reflect ambiguous terminology, differences in annotation specificity, or potentially incorrect reference assignments. Hierarchy-aware evaluation can help characterize predictions related to the reference concept, but ancestor and descendant predictions should not automatically be considered equivalent to exact normalization. In particular, descendant predictions may introduce specificity not supported by the original mention. Strict concept-level accuracy therefore remains the primary evaluation criterion.

Third, the performance of pretrained models may depend on the terminology and annotations represented in their training resources. Although SapBERT performed competitively across the evaluated ontologies, this study does not isolate the influence of UMLS terminology coverage from other factors affecting retrieval. Similarly, the benefits of ontology-specific adaptation were examined only for cell-type normalization, and further evaluation would be needed to determine whether comparable improvements extend to other biological ontologies.

Finally, the present evaluation addresses entity normalization rather than the complete process of extracting and integrating biological relationships from scientific literature. Although the workflow was motivated by the requirements of rare disease knowledge integration, its performance was assessed using existing normalization benchmarks. Its contribution to downstream information extraction, including the accuracy and coverage of extracted relationships, remains to be evaluated within the intended application.

## Conclusions

This study provides a systematic evaluation of a modular biomedical entity normalization workflow across heterogeneous biological ontologies relevant to rare disease knowledge integration. The results show that normalization performance depends on both candidate retrieval and concept selection, with substantial variation across entity types and evaluation benchmarks. Combining lexical and semantic retrieval can improve candidate coverage in some settings, while LLM-based reranking provides an effective selection strategy. The improvements obtained with SapBERT-cell further illustrate the potential value of incorporating ontology-adapted retrieval models when suitable resources are available.

The workflow offers a common approach for normalizing different biological entity types without requiring separate ontology-specific training, while retaining the flexibility to incorporate specialized components. It provides a methodological foundation for extracting and integrating biological knowledge from rare disease literature, with potential applicability to other biomedical information-extraction tasks involving the same ontologies.

## Methods

### Study design

We developed a general-purpose biomedical entity normalization workflow and evaluated its performance across six biological entity types: phenotypes, cell types, anatomical structures, biological processes, molecular functions, and cellular components. The target terminology resources were the Human Phenotype Ontology (HPO), Cell Ontology (CL), Uberon, Gene Ontology (GO), and Unified Medical Language System (UMLS). The workflow comprises three stages:

1. **Exact matching.** Input mentions were compared with preferred labels and synonyms from the target terminology after applying a common text normalization procedure (detailed below). All available synonym categories (exact, related, narrow, and broad, where applicable) were considered. Mentions matching a preferred label or synonym were resolved at this stage and excluded from subsequent candidate retrieval and reranking.
2. **Candidate retrieval.** For mentions without an exact match, candidate ontology concepts were retrieved and ranked using four complementary methods: BM25, fuzzy string matching, character n-gram similarity, and SapBERT. Each method retrieved all unique candidate concepts with ranks up to 100, using standard competition ranking for tied scores (i.e., tied candidates received the same rank, with subsequent ranks adjusted accordingly). Consequently, more than 100 candidates could be retrieved when ties occurred at the cutoff. Candidate lists from individual methods were also merged and deduplicated to evaluate the complementarity of lexical and semantic retrieval.
3. **Candidate selection.** Retrieved candidates were ranked or selected using three general-purpose strategies: LLM-based selection, Borda count, and reciprocal rank fusion (RRF). These methods were applied without ontology-specific training or fine-tuning.

SapBERT was selected as the semantic retrieval method because it was specifically trained to align synonymous biomedical concepts in a shared embedding space, making it suitable for biomedical entity normalization (Liu et al., 2020). The study focused on comparing complementary retrieval strategies rather than benchmarking alternative semantic representation models.

### Evaluation datasets

We initially considered four biomedical annotation benchmarks: MedMentions for UMLS, CellLink for the Cell Ontology (CL), PhenoNormBench for the Human Phenotype Ontology (HPO), and CRAFT for CL, Uberon, and the biological process (BP), molecular function (MF), and cellular component (CC) branches of the Gene Ontology (GO). Following the analysis of exact matching and the availability of non-exact mentions, CRAFT was retained for GO normalization, while CellLink and a separately constructed FMA–Uberon dataset were used for the non-exact normalization evaluations of cell types and anatomical structures, respectively.

For MedMentions, we selected mentions corresponding to six biological entity types based on their UMLS semantic types: cell types (T025); phenotypes (T046 and T048); anatomical structures (T023, T024, T029, T030, and T022, excluding T026, T028, and T021); biological processes (T042 and T043); molecular functions (T044); and cellular components (T026). These semantic-type groupings do not necessarily correspond exactly to the scope of the individual target ontologies.

For anatomical structure normalization against Uberon, we selected 18,812 UMLS terms associated with the Foundational Model of Anatomy (FMA) and mapped their FMA identifiers to Uberon using ontology cross-references, obtaining 6,046 distinct reference concepts. Unlike the other benchmarks, this dataset evaluates normalization of terminology entries rather than mentions extracted from biomedical text.

### Ontology and terminology preprocessing

We obtained the Cell Ontology (CL; June 8, 2026), Uberon (June 25, 2023), Human Phenotype Ontology (HPO; June 23, 2026), and Gene Ontology (GO; June 15, 2026) in OWL format. For each ontology, we extracted concept identifiers, preferred labels, and synonyms, including exact, narrow, broad, and related synonyms, as well as hierarchical relationships between concepts.

For the Unified Medical Language System (UMLS), we used the 2026AA release. Concept unique identifiers (CUIs) and their associated terms were extracted from MRCONSO.RRF, while semantic type assignments were obtained from MRSTY.RRF.

### Identification of exact lexical matches

Input mentions and ontology terms were standardized using Unicode normalization, case folding, whitespace normalization, and punctuation standardization. Each standardized mention was then compared against the preferred labels and synonyms of the target ontology.

When a mention matched terms associated with multiple concepts, matches were prioritized according to the following order: preferred labels, exact synonyms, broad synonyms, narrow synonyms, and related synonyms. For UMLS, mentions were compared against all terms associated with each CUI.

Exact-match precision was calculated as the proportion of exactly matched mentions for which the assigned concept agreed with at least one reference annotation, when multiple reference concepts were provided. We additionally performed a post-hoc hierarchy-aware evaluation of predictions that did not match any reference identifier. Two progressively broader criteria were considered: (i) accepting predictions corresponding to an ancestor of at least one reference concept, and (ii) accepting predictions corresponding to either an ancestor or a descendant of at least one reference concept. Hierarchical relationships were evaluated transitively, including indirect ancestors and descendants.

### Candidate retrieval for non-exact terms

Mentions without an exact match were processed using four complementary candidate retrieval methods: three lexical approaches (BM25, fuzzy string matching, and character n-gram similarity) and one semantic approach (SapBERT).

The lexical methods capture different aspects of similarity between input mentions and ontology terms. BM25 ranks terms according to token overlap, weighted by their informativeness within the terminology. Fuzzy matching identifies terms with similar character sequences, accommodating minor spelling differences and typographical variations, while character n-gram similarity captures partial lexical overlap through shared character subsequences. For BM25, input mentions were lemmatized using scispaCy, following preliminary experiments indicating improved retrieval performance with this preprocessing step.

Semantic retrieval was performed using SapBERT (Liu et al., 2020), a biomedical representation model trained to align synonymous concepts from UMLS in a shared embedding space. Input mentions and ontology terms were represented as dense embeddings, and candidate ontology terms were retrieved according to their semantic similarity. The pretrained model was applied to all target ontologies without additional fine-tuning.

For all four retrieval methods, similarity scores were calculated independently between each input mention and the textual representations of the target concepts, including preferred labels and synonyms. Each concept was assigned the maximum score obtained across its textual representations, and unique concepts were subsequently ranked in descending order of score. We retained all concepts with ranks up to 100, using standard competition ranking for tied scores: concepts with identical scores received the same rank, and subsequent ranks accounted for the number of preceding concepts. Consequently, candidate lists could contain more than 100 concepts when ties occurred at the cutoff. We evaluated both individual retrieval methods and candidate sets obtained by combining and deduplicating their results.

Candidate retrieval performance was assessed using Recall@k, defined as the proportion of non-exact mentions for which at least one reference concept was retrieved with a rank of (k) or better. For mentions annotated with multiple reference concepts, retrieving any one of them was considered successful.

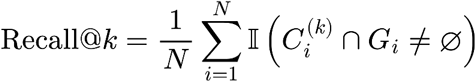

where (*N*) is the total number of unique non-exact mentions in the evaluation set, *C_i_^(k)^* denotes the set of candidate concepts retrieved for mention (*i*) with a rank of (*k*) or better, and (*G_i_*) represents the set of reference concepts annotated for that mention. The indicator function (Ⅱ(·)) takes a value of 1 if the intersection between the retrieved candidates and the reference concepts is non-empty (i.e., at least one reference concept is retrieved within the top (*k*)), and 0 otherwise.

### Candidate reranking and evaluation of selection accuracy

Following candidate retrieval, we evaluated three methods for selecting the concept most likely to correspond to each input mention: large language model (LLM)-based reranking, reciprocal rank fusion (RRF), and Borda rank aggregation.

#### LLM-based reranking

We used the deepseek-ai/DeepSeek-V4-Flash model to select the most appropriate concept from the retrieved candidates. The model was provided with the input mention and the preferred labels and synonyms of the candidate concepts and was prompted to identify the best match. The complete system prompt is provided in Supplementary Note S1.

#### Reciprocal rank fusion (RRF)

We used RRF to combine the rankings produced by the four retrieval methods, assigning higher scores to concepts appearing at higher positions in the individual rankings. For each candidate concept (c), the aggregated score was calculated as:

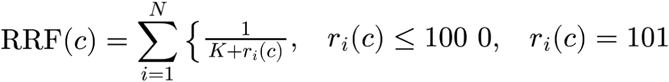

where (*N* = 4) is the number of retrieval methods, (*r_i_*(*c*)) is the rank assigned to candidate (*c*) by method (*i*), and (*K* = 60) is a smoothing constant. Candidates were ranked in descending order of their aggregated scores.

#### Borda rank aggregation

We used a rank-sum approach based on the Borda count method to combine the rankings produced by the four retrieval methods:

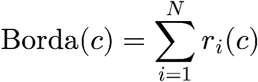

where (*N* = 4) and (*r_i_*(*c*)) is the rank assigned to candidate (*c*) by retrieval method (i). Candidates were ranked in ascending order of their aggregated scores.

For both rank aggregation methods, candidate retrieval was limited to concepts with ranks up to 100 in each individual method. Concepts absent from a method’s retrieved list were assigned a rank of 101, representing candidates beyond the retrieval cutoff. This rank contributed zero to RRF and a value of 101 to the Borda rank sum. Tied retrieval scores retained their shared ranks, and ties could also occur after rank aggregation.

To investigate the effect of candidate-set size on reranking performance, we evaluated candidate sets constructed by merging and deduplicating the top-ranked concepts retrieved by the individual methods at different rank cutoffs (k). Because the number of unique candidates could vary across mentions, each experimental setting was characterized by the median number of unique candidates per mention. LLM-based reranking was evaluated only for the candidate-set sizes considered in the experiments, rather than for the full retrieval lists extending to rank 100.

Reranking performance was evaluated using RerankingAccuracy@k, defined as the proportion of mentions for which the reranking method selected at least one correct reference concept, considering only mentions for which at least one reference concept was present in the candidate set:

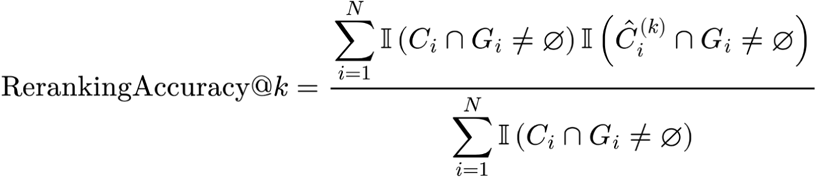

where (*N*) is the total number of unique non-exact mentions, (*C_i_*) denotes the candidate set retrieved for mention (*i*), (*G_i_*) represents its set of reference concepts, and (*Ĉ_i_*^(*k*)^) denotes the top-k concepts selected by the reranking method. The indicator function (Ⅱ(·)) returns 1 when its argument is true and 0 otherwise. The denominator therefore counts only eligible mentions, for which at least one reference concept was present in the original candidate set, whereas the numerator counts those eligible mentions for which at least one reference concept was retained among the top-k reranked candidates.

For mentions annotated with multiple reference concepts, selecting any one of them was considered correct. Similarly, when multiple candidates shared the highest reranking score, the prediction was considered correct if at least one of these candidates matched a reference concept. Thus, RerankingAccuracy@k measures candidate selection performance conditional on successful retrieval, excluding mentions for which no reference concept was retrieved.

### Evaluation of overall normalization performance

After evaluating candidate retrieval and reranking separately using Recall@k and RerankingAccuracy@k, respectively, we assessed the performance of the complete normalization pipeline using OverallAccuracy@k. Unlike RerankingAccuracy@k, which evaluates candidate selection only for mentions where at least one reference concept was retrieved, OverallAccuracy@k considers all evaluated mentions. It is defined as the proportion of mentions for which the reranking method selects at least one correct reference concept from the candidate set.

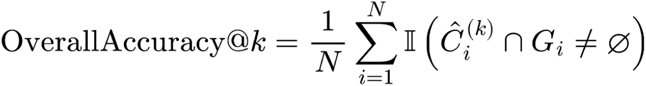

where (*N*) is the total number of unique mentions evaluated, (*G_i_*) represents the set of reference concepts for mention (*i*), and (*Ĉ_i_*^(*k*)^) denotes the top-k concepts selected by the reranking method. The indicator function (Ⅱ(·)) returns 1 when at least one reference concept is present among the top-k selected candidates and 0 otherwise. This metric includes all evaluated mentions in the denominator, including those for which no reference concept was retrieved during candidate generation.

In addition to strict identifier matching, we performed a hierarchy-aware evaluation of all performance metrics. Under this criterion, a prediction was also considered correct if the selected concept was an ancestor or descendant of a reference concept in the corresponding ontology, including indirect hierarchical relationships. This analysis accounts for predictions that differ from the reference annotation in their level of specificity.

### Characterization of retrieval difficulty

We investigated factors associated with candidate retrieval performance using the CellLink validation dataset. Retrieval difficulty was quantified by the best (lowest) rank assigned to any reference concept for each mention and retrieval method. Mentions for which no reference concept was retrieved within the rank cutoff of 100 were assigned a rank of 101.

We first used Spearman’s rank correlation coefficient to assess associations between retrieval rank and three characteristics: mention length, measured by the number of characters and tokens, and the number of synonyms associated with the reference concept in the Cell Ontology (CL). Correlations were calculated separately for each retrieval method.

We then examined whether mentions containing lists of biological entities differed in retrieval difficulty from those referring to a single entity. The distributions of reference concept ranks were compared using the two-sample Kolmogorov–Smirnov (KS) test.

Finally, we investigated whether retrieval performance varied across categories defined by the CL hierarchy. For each selected parent concept, we compared the distributions of reference concept ranks between mentions annotated with concepts belonging to its descendant hierarchy (including indirect descendants) and all other mentions, using the two-sample KS test.

### Comparison with a specialized cell-type normalization system

We compared our general-purpose normalization pipeline with CellExLink (Nabijiang & Shahriyari, 2026), a specialized biomedical text-mining system for extracting and normalizing cell-type mentions to the CL. To evaluate the contribution of ontology-specific model adaptation to candidate retrieval, we compared CellExLink retrieval method with two semantic embedding models within our pipeline: the original pretrained SapBERT and SapBERT-cell, a version fine-tuned by the CellExLink authors using CL terminology and annotated cell-type mentions. All three methods were evaluated on the CellLink test subset used in the CellExLink study, which was not included in the fine-tuning data for SapBERT-cell. Candidate retrieval performance was assessed using Recall@k. In addition, to compare the overall performance of our general-purpose normalization framework with the complete CellExLink pipeline, we calculated OverallAccuracy@k for both systems using the candidates retrieved and reranked from the same CellLink test subset.

## Code availability

The code and scripts developed for this study, including the entity normalization pipeline and evaluation scripts are publicly available on GitHub at https://github.com/KnownRD/entity-normalization

