## Supplementary material for "A modular workflow for biomedical entity normalization across heterogeneous biological ontologies"

**Table S1: Number of unique entities across benchmark.** Values in parentheses indicate non-exact unique entities

| | MedMentions | CellLink | CRAFT | PhenoNormBench | FMA-UMLS |
|---|---|---|---|---|---|
| Cell types | 1,428 (1,021) | 5,156 (4,672) | 790 (637) | - | - |
| Phenotypes | 1,310 (657) | - | - | 6,136 (4,483) | - |
| Anatomical structures | 1,890 (386) | - | 1,037 (3) | - | 18,812 (1,913) |
| Biol. process | 1,508 (1,009) | - | 1,344 (945) | - | - |
| Mol. func. | 1,107 (816) | - | 539 (499) | - | - |
| Cell comp. | 541 (319) | - | 585 (441) | - | - |

**Table S2. Precision of exact terms.** Number of distinct exact-match terms and exact-match precision for MedMentions benchmark.

| | **Number exact terms** | **Precision** |
|---|---|---|
| MedMentions cell types | 407 | 0.926 |
| MedMentions anatomy | 1.504 | 0.862 |
| MedMentions phenotypes | 653 | 0.916 |
| MedMentions biological processes | 499 | 0.962 |
| MedMentions molecular functions | 291 | 0.914 |
| MedMentions cellular components | 222 | 0.946 |

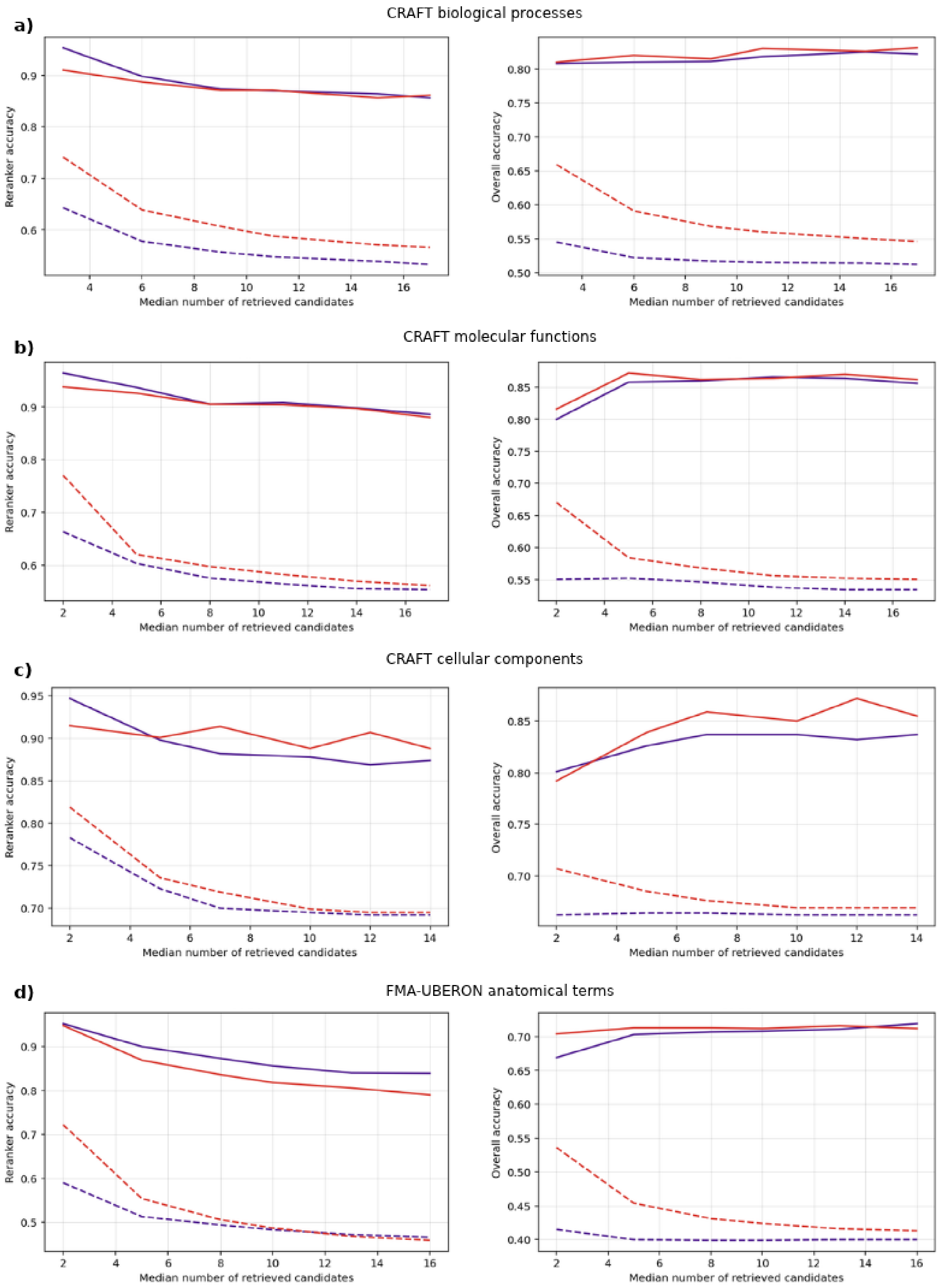


**Fig S1. Candidate selection performance across specialized biomedical ontologies**. Reranker and Overall Accuracy@k across candidate-set sizes. These metrics are calculated using LLM-based and Borda count rerankers in two retrieval scenarios: combined candidates, consisting of candidates retrieved by lexical and SapBERT methods and only SapBERT candidates. The representation includes both metrics in different benchmarks Gene Ontology (GO) annotations in CRAFT for (a) biological processes, (b) molecular functions and (c) cellular components, and (c) Uberon in FMA–UBERON.

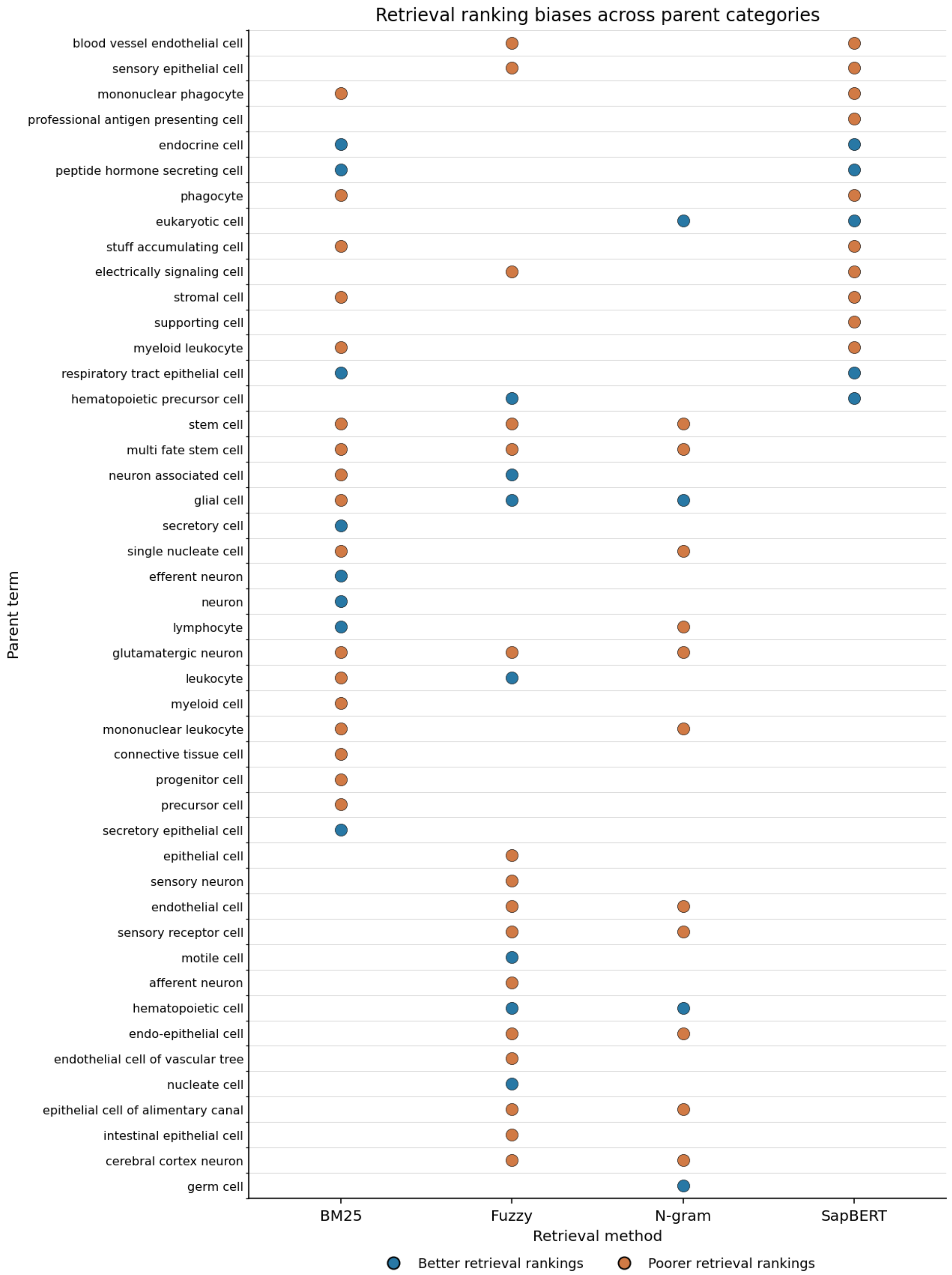


**Fig S2. Retrieval ranking biases across parent categories.** Dot plot showing biases in the retrieval rankings of gold-standard CellLink concepts according to their parent categories across BM25, fuzzy matching, character n-gram, and SapBERT retrieval methods. Blue dots indicate a significant association with better retrieval rankings, whereas orange dots indicate a significant association with poorer retrieval rankings (KS test, $p < 0.01$). Categories without significant associations are represented without a dot.

**Supplementary Note S1. System prompt for LLM-based reranking.**

SYSTEM_PROMPT = """
*You are a professional data annotator and curator.*

*Given a query and candidate IDs with their associated ontology terms,*
*select the candidate ID that most specifically and biologically correctly represents*
*the query. Prefer a candidate whose ontology term denotes the same biological entity*
*as the query.*

*You MUST provide an ANSWER from among the candidates*

*Return ONLY valid JSON:*
*{"ID": "candidate"}*

*Do not provide explanations.*
"""
